# SEX DIFFERENCES IN BLOOD PRESSURE RESPONSE TO CONTINUOUS ANG II INFUSION: ARE ONLY SEX HORMONES TO BLAME?

**DOI:** 10.1101/2021.10.15.464458

**Authors:** Florencia Dadam, Andrea Godino, Laura Vivas, Ximena E. Caeiro

## Abstract

To investigate the involvement of the sex chromosome complement (SCC), organizational and activational hormonal effects in changes in mean arterial pressure during acute Ang II infusion, we used gonadectomized (GDX) mice of the “four core genotypes” model, which dissociates the effect of gonadal sex and SCC, allowing comparisons of sexually dimorphic traits between XX and XY females as well as XX and XY males. Additionally, β-estradiol and testosterone propionate (2ug/g) were daily injected for 4 days to evaluate activational hormonal effects.

Statistical analysis of the changes in mean arterial pressure revealed an interaction of SCC, organizational and activational hormonal effects during Ang II infusion {F(7,39=2,60 p<0.01)}. Our results indicate that, in absence of activational hormonal effects, interaction between the SCC and organizational hormonal action differentially modulates changes in arterial pressure. In GDX mice without hormone replacement, Ang II infusion resulted in an increase in mean arterial pressure in XX-male, XX-female and XY-female mice, while no changes were observed in XY-male mice. Furthermore, β-estradiol replacement (GDX+E2 group) resulted in a decrease in blood pressure in XX-males, XX-females and XY-females (indicating an activational β-estradiol effect), while no changes were observed in the XY-male group. Moreover, testosterone propionate replacement (GDX+TP group) showed a greater increase in blood pressure in XY-male mice than in XX-males and XX-females, demonstrating an activational hormonal effect of testosterone in XY-male mice.

Our data isolates and highlights the contribution and interaction of SCC, activational and organizational hormonal effects in sex differences in Ang II blood pressure regulation.

## INTRODUCTION

Although awareness of sex differences in cardiovascular disease is increasing, much of what we know about blood pressure regulation has been derived from studies in males. However, principles learned in male models do not necessarily apply to females, and it is therefore important to understand in more detail the physiological basis of sex differences. Sexual dimorphism in blood pressure regulation under physiological and pathophysiological states has long been recognized in both clinical and experimental studies (Calhoun et al., 1995; Fischer et al., 2002; Reckelhoff et al., 2000). The renin angiotensin system (RAS) induces both hormonal and paracrine effects, modulating blood pressure regulation, among others. Ang II acts physiologically by binding to two G protein-coupled receptor subtypes, the angiotensin II type-1 receptor (AT1R) and angiotensin II type-2 receptor (AT2R). AT1R activation promotes vasoconstriction, antinatriuresis, and sympathetic nervous activation, as well as inflammatory, thrombotic, proliferative, and fibrotic processes (de Gasparo et al., 2000), while binding to AT2R evokes vasodilatation, natriuresis, anti-inflammation, and anti-proliferation, generally opposing the AT1R-mediated responses (Verdonk et al., 2012). Furthermore, Ang II is degraded to Ang (1-7) by ACE2 which, through binding to MasR, triggers anti-inflammatory, anti-fibrotic, and antiproliferative actions. Thus, RAS includes two counter arms, the vasoconstrictor arm (Ang-converting enzyme (ACE)/AngII/AT1R), and the protective arm (ACE2/Ang (1-7)/Mas receptor (MasR) (Liu et al., 2019).

Clinical and basic findings have demonstrated major sex differences in the way males and females respond to stimulation and inhibition of the RAS under physiological and pathophysiological circumstances (Brown et al., 2012; Sullivan, 2008; Sullivan et al., 2010; Xue et al., 2005). But why are there male and female differences in the way RAS modulates blood pressure? Exposure to sex steroids during critical periods of development can induce organizational (long-lasting or permanent) effects on sexually dimorphic traits. Sex steroids can also impart (temporary or reversible) activational effects at different times of life (during neonatal and peripubertal development as well as in adulthood), causing many known sex differences in phenotypes (Arnold and Gorski, 1984; McCarthy et al., 2012; Morris et al., 2004). For a long time, the hormonal organizational-activational dichotomy was applied to the understanding of most sex differences, and hormones were the only factors discussed as proximate signals causing sex differences. However, males and females differ not only in their sex (males are born with testes- and females with ovaries-hormonal factors) but also carry different sex chromosome complements (SCC: XY and XX respectively), and thus are influenced throughout life by different genomes.

With regard to activational hormonal effects, studies have demonstrated that chronic treatment with estradiol attenuates hypertension in intact males and ovariectomized females, while administration of testosterone increases hypertension in gonadectomized males (Crofton and Share, 1997). Testosterone regulates (increases) angiotensinogen production, renin activity, and the expression of the AT1-r receptor, and downregulates AT2 receptor expression (Mishra et al., 2016), favoring in this case the vasoconstrictor effect of RAS, and these effects are significantly diminished after castration in males (Sampson et al., 2012). On the other hand, estrogens are known to increase angiotensinogen, AT2 density, and eNOS, while they decrease not only renin but also AT1 receptor density. Furthermore, estrogen reposition in OVX females reveals a downregulation of angiotensin-converting enzyme (ACE) transcript levels (Hilliard et al., 2013; Gallagher et al., 1999).

With regard to the contribution of the sex chromosome complement to physiological sex-based differences in the regulation of RAS in blood pressure regulation, it is important to note that two genes of the protective RAS arm, AT2 and ACE2, are located on the X-chromosome [25]. Exciting new data indicate that some genes escape X-inactivation and are expressed from both the “active” and the “inactive” X chromosome, which may cause functional sex differences intrinsic to male (XY) and female (XX) cells, potentially contributing to sex differences in traits (sex-biased genes) (Carrel and Willard, 2005; Wolstenholme et al., 2013; Yang et al., 2006). Previous evidence demonstrates a modulatory effect of SCC on RAS receptor expression (brain and renal) (Dadam et al., 2017), as well as in the Ang II sexually dimorphic bradycardic baroreflex (Caeiro et al., 2011) and hypertensive responses (Ji et al., 2010). However, in all these studies, SCC and organizational hormonal effects were evaluated in absence of activational hormonal effects.

Many studies indicate that sex differences in ACE activity and AT1R/AT2R expression/sensitivity may account for some of the Ang II-related sex differences associated with the vasoconstrictor/vasodilator balance of the RAS (Sullivan, 2008; Silva-Antonialli et al., 2004). In intact animals, enhancement of the vasodilation component of RAS in females has also been described, as the infusion of low doses of Ang II potentially biases females in the direction of vasodilation and males toward vasoconstriction (Sampson et al., 2008). But are these differences among males and females due only to hormonal activation and/or organizational effects?

In view of these and previous studies that demonstrate an important activational hormonal effect, in the present study we sought to evaluate the involvement of not only SCC and the organizational hormonal effect but also the activational hormonal role of testosterone and estradiol in sexual dimorphism in changes in mean arterial pressure (MAP) during continuous acute Ang II infusion.

## METHODS

### Animals

Transgenic mice of the “four core genotypes” mouse model were used in the experiments. MF1 transgenic mice, kindly provided by Dr. Paul Burgoyne from the Medical Research Council National Institute for Medical Research, UK, were born and reared in the breeding facilities at the Instituto Ferreyra (Córdoba, Argentina). This mouse model combines a deletion of the testis-determining Sry gene from the Y chromosome (Y_) and the subsequent insertion of an Sry transgene into an autosome. Sry gene deletion in XY mice (XY_) yields a female phenotype (ovaries). When the Sry transgene is inserted into an autosome of these mice, they have testes and are fully fertile (XY_Sry). The Y_ chromosome and the Sry transgene segregate independently, and thus four types of offspring are produced by breeding XY_Sry males with XX females: XX and XY_ females (without Sry on the Y chromosome) and XX-Sry and XY_Sry male mice (both with Sry in an autosome). All individuals possessing the Sry transgene develop testes and have a male external phenotype regardless of their SCC, while individuals lacking the transgene have ovaries and external female secondary sex characteristics.

Male and female are defined here according to the gonadal phenotype. Throughout the text, we will refer to XX and XY-as XX-female and XY-female, and to XX-Sry and XY_Sry as XX-male and XY-male mice respectively. By comparing these genotypes, it is possible to segregate the role of a) SCC (comparing mice with the same gonadal type but with different SCC: XX vs. XY), b) gonadal sex (males vs. females regardless of SCC), and c) the interaction of SCC and gonadal sex (Fig. 1). Furthermore, when activational hormonal effects were also to be tested, we used gonadectomized mice with hormone replacement, thus evaluating the involvement of SCC, organizational and activational hormonal effects in sexually dimorphic phenotypes (Fig. 1).

**Fig. 1.**
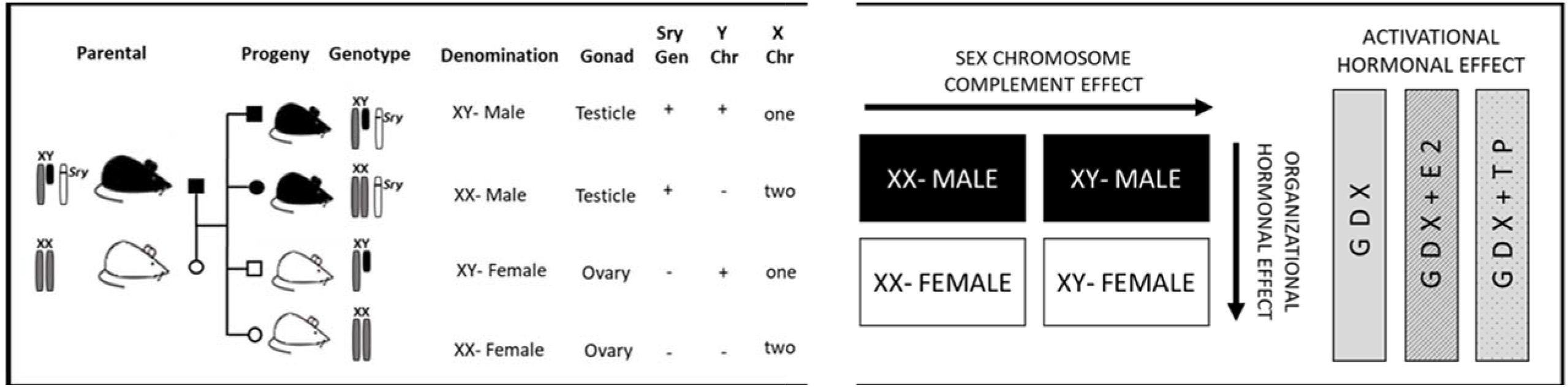
Schematic representation of four core genotypes mouse model (FCG). Left panel: Four types of offspring are produced by breeding XY_Sry males to XX females: XX and XY_females (without Sry on the Y chromosome) and XXSry and XY_Sry male mice (both with Sry in an autosome). All of the individuals possessing the Sry transgene develop testes and have a male external phenotype, regardless of their SCC, whereas individuals lacking the transgene have ovaries and external female secondary sex characteristics. XX and XY_ females are referred as XX and XY females and XXSry and XY_Sry male mice as XX and XY male mice, respectively. Right panel: comparisons between the genotypes in the FCG model to discriminate between the SCC effect, (comparing XY-females vs. XX-females and XY-males vs. XX-males) the gonadal sex (comparing males vs. females), and activational hormonal effect (comparing GDX, GDX+E2 and GDX+TP).

Genotyping was performed as previously described (Caeiro et al., 2011). All experimental protocols were approved by the appropriate animal care and use committees at our institute (001/2019), following the National Institutes of Health guidelines for the care and use of laboratory animals.

### Gonadectomy – Surgical procedures

Adult mice were anesthetized with ketamine/xylazine and a bilateral incision was made, in the scrotum region for male mice and just below the rib cage in the female mice, to be able to perform bilateral gonadectomy. Then the vascular supply was ligated, the gonads (either testes or ovaries) were removed, and the muscle layer and incisions were sutured in place.

### Experimental groups and designs

Mice of the four genotypes (aged 45/50 days old) were gonadectomized to remove any activational effect of sex hormones that might mask the modulatory action of SCC and the organizational hormonal effects. As shown in Figure 2, after 12 days of gonadectomy, animals were divided into three groups: a) without hormone replacement (GDX group, to evaluate SCC and organizational hormonal effects), b) with β-estradiol replacement (GDX+E2 group, to evaluate SCC, organizational and activational β-estradiol effects), and c) with testosterone propionate replacement (GDX+TP group, to dissociate SCC, organizational and activational testosterone propionate effects). The groups with hormone replacement were treated for 4 consecutive days with daily subcutaneous injections of β-estradiol or testosterone propionate (2ug/gpc/day).

**Fig. 2.**
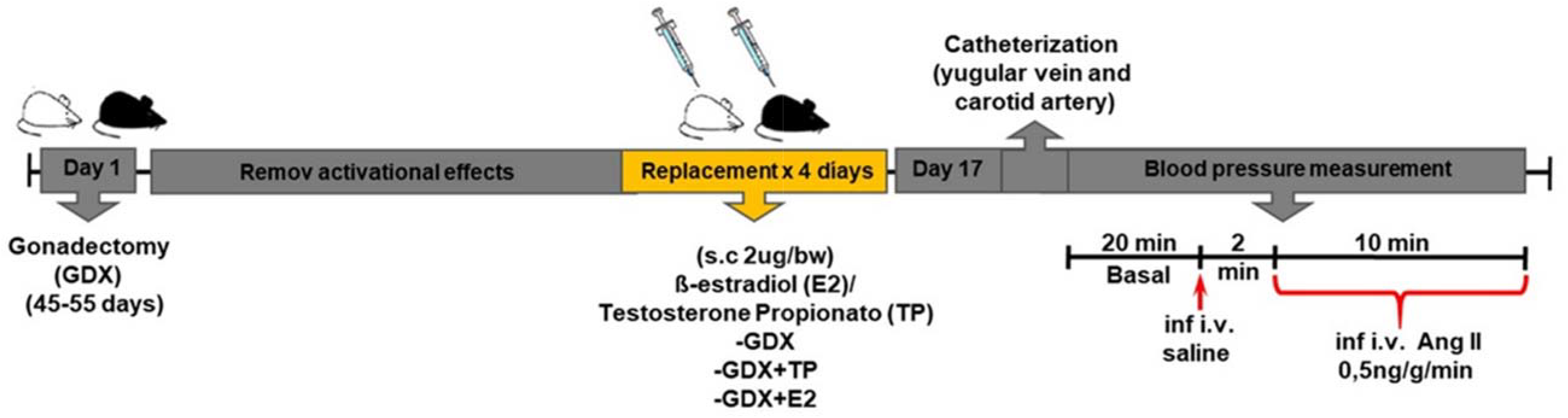
Schematic representation of the experimental design. Twelve days after gonadectomy, the animals were assigned to three experimental groups: without hormonal replacement (GDX), with β-estradiol replacement (GDX + E2) and with testosterone propionate replacement (GDX + TP). These last two groups were treated for 4 consecutive days with daily subcutaneous replacement of the corresponding hormone (2 ug / gpc / day). After 17 days, mice underwent surgery for cannulation of the carotid artery and jugular vein. Baseline parameters were taken during the first 20 minutes and then a continuous infusion of Ang II was performed during the next 10 minutes (Ang II, Sigma, 250 μg / mL; infusion rate of 0.003 ml / min). Blood pressure and heart rate were recorded.

Seventeen days after gonadectomy, mice of three groups (GDX/GDX+E2/GDX+TP) were anesthetized with urethane and were subjected to cannulation of the carotid artery and jugular vein (Caeiro et al., 2011). After surgery, the arterial catheter was connected to the blood pressure transducer coupled to a data acquisition system (Power Laboratory, ADI instruments), and the venous catheter was coupled to a pump for small volume infusion. After 20 minutes of baseline recording, vehicle solution was infused and, two minutes later, continuous infusion of Ang II was conducted for the following 10 minutes (Ang II, Sigma, 250 μg/mL; infusion rate of 0.003ml/min) and blood pressure and heart rate were recorded. The experimental protocol was carried out between 9:00 AM and 4:00 PM.

## STATISTICAL ANALYSIS

Data of changes in mean arterial pressure were subjected to a 3-way mixed ANOVA with repeated measures; organizational sex (male/female), activational hormonal effect (GDX/GDX+E2/GDX+TP), and SCC (XY/XX) factors were considered as independent factors. The loci of significant interactions or significant main effects were further analyzed using the Tukey test (type I error probability was set at 0.05). Data of changes in blood pressure in each genotype were subjected to a 1-way ANOVA with repeated measures, and treatment factor (GDX/GDX+E2/GDX+TP) was considered as an independent factor. Results were expressed as group mean (M) ± standard error (SE).

## RESULTS

The statistical analysis of changes in blood pressure in response to Ang II infusion reveals an interaction of SCC, organizational-sex and activational-hormonal effect {F(7,39=2,60 p<0.01)}. In control groups (gonadectomized mice without hormone replacement), Ang II infusion resulted in an increase in mean arterial pressure in XX-male, XX-female and XY-female mice (XX-Male/GDX, XX-Female/GDX, XY-Female/GDX), while no changes were observed in XY-male mice (XY-Male/GDX) (Fig. 3A).

**Fig. 3.**
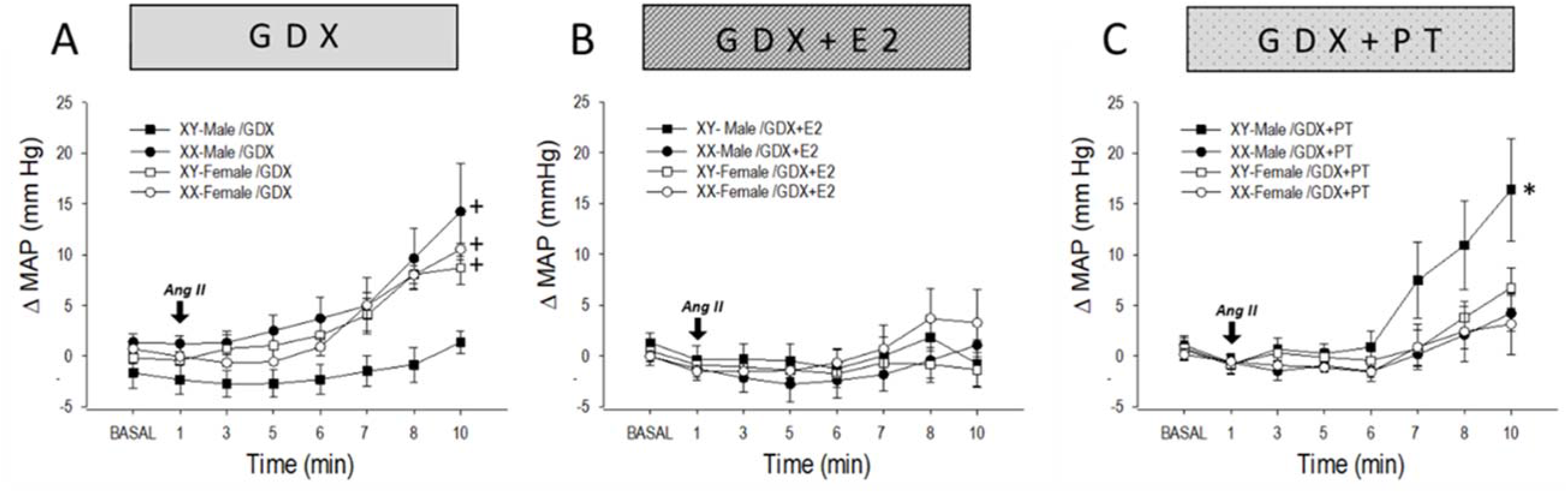
Changes in mean arterial pressure (ΔMAP) during continuous Ang II infusion (Panel A-C) and during 10 minutes of Ang II infusion in gonadectomized mice without hormone replacement (GDX group, **Panel A**), in gonadectomized mice with β-estradiol replacement (GDX+E2, **Panel B**) and in gonadectomized mice with testosterone propionate replacement (GDX+TP, **Panel C**). n=6-9/group. *p<0.01: XY-Male/GDX+TP vs. XX-Male/GDX+TP, XX-Females/GDX+TP, XY-Male/GDX+E2, XX-Male/GDX+E2, XX-Females/GDX+E2, XY-Females/GDX+E2 and XY-Male/GDX. +p<0.01: XX-Male/GDX, XX-Female/GDX and XY-Female/GDX vs. XY-Male/GDX, XY-Male/GDX+E2, XX-Male/GDX+E2, XX-Female/GDX+E2, and XY-Female/GDX+E2.

Furthermore, β-estradiol replacement (E2 group) resulted in a decrease in blood pressure in XX-male, XX-female and XY-female mice (XX-Male/GDX+E2, XX-Female/GDX+E2 XY-Female/GDX+E2) compared to the same experimental groups without hormone replacement (indicating an activational effect of β-estradiol). However, no changes in mean arterial pressure were observed among XY-male mice, with or without β-estradiol replacement (XY-Male/GDX vs. XY-Male/GDX+E2) (Fig. 3A and B).

The statistical analysis of experimental groups that received testosterone propionate replacement showed a greater increase in blood pressure in XY-Male/GDX+TP mice than in XX-male and XX-female/GDX+TP groups, demonstrating an activational hormonal effect of testosterone in XY-male mice (Fig. 3C).

When changes in mean arterial pressure in each genotype were analyzed (Fig. 4) under the different experimental conditions (without hormone replacement, and with estradiol and propionate testosterone replacement, respectively), the results demonstrate in the XY-Male genotype that GDX+TP mice showed an increase in blood pressure compared to XY-Male/GDX or XY-Male/GDX+E2 mice (in which no increase in mean arterial pressure was observed), thus demonstrating the pressor activational hormonal effect of testosterone in XY-Male mice (Fig. 4, panel A).

**Fig. 4.**
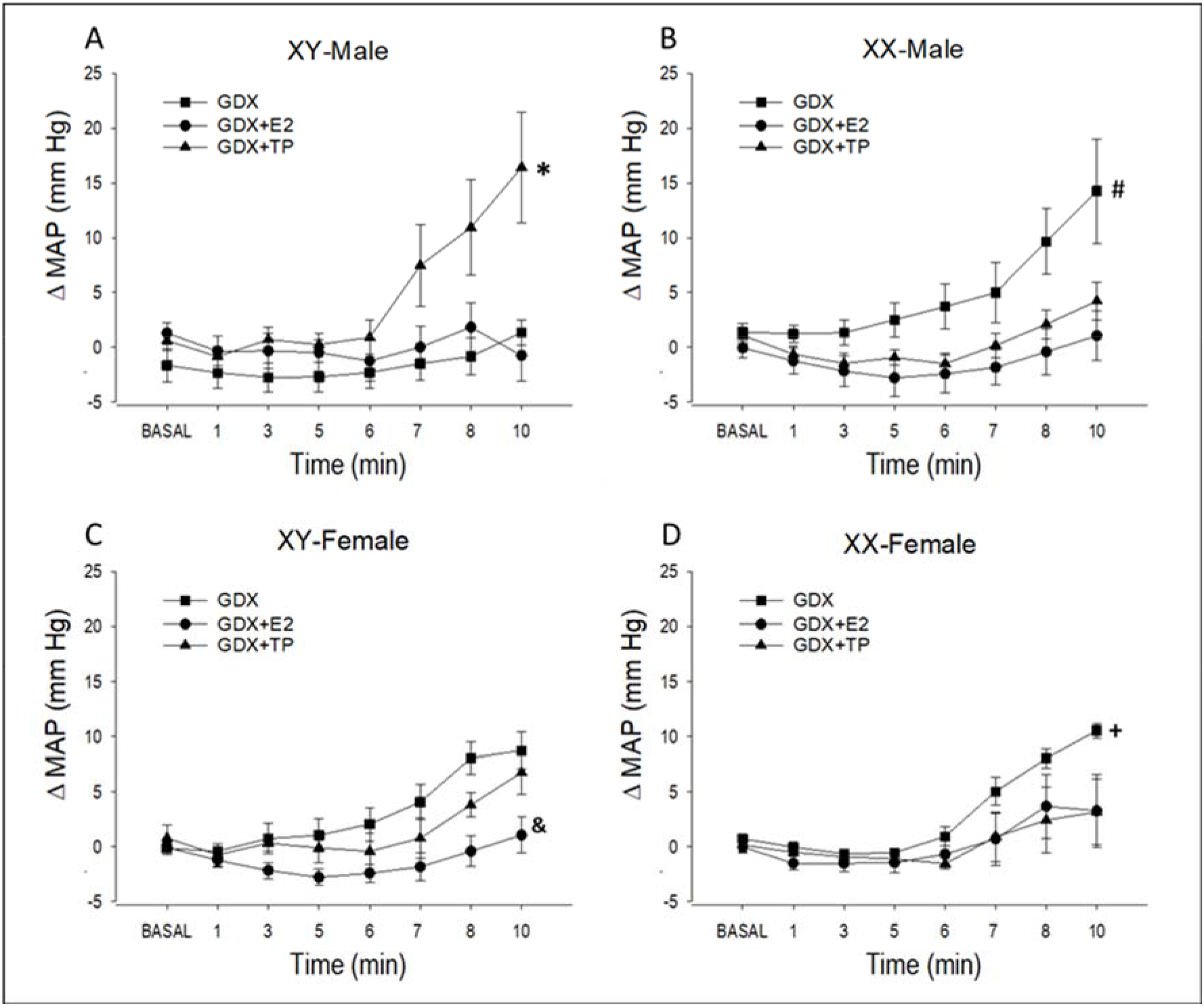
Changes in mean arterial pressure (ΔMAP) induced by continuous infusion of Ang II. (Ang II, Sigma, 250 μg/mL; infusion rate 0.003ml/min) in: A) male-XY, B) male-XX, C) female-XY and D) female-XX mice under different experimental situations: gonadectomized without hormone replacement (GDX, square), gonadectomized with β-estradiol replacement (GDX+E2, circle) and with testosterone propionate replacement (GDX+TP, triangle). n=6-9/group. *p<0.001: XY-Male/GDX+TP vs. XY-Male/GDX and XY-Male/GDX+E2. #p<0.01: XX-Male/GDX vs. XX-Male/GDX+E2 and XX-Male/GDX+TP. &p<0.001: XY-Female/GDX+E2 vs. XY-Female/GDX and XY-Female/GDX+TP. +p<0.05: XX-Female/GDX vs. XX-Female/GDX+E2.

Moreover, as shown in panel B, gonadectomized XX-male mice without hormone replacement (XX-male/GDX) showed an increase in blood pressure when Ang II was infused and, with estradiol replacement (XX-male/GDX+E2), a depressor activational effect was observed. In line with what was observed in the group of XX-male mice, in XY- and XX-gonadectomized female mice without hormone replacement (XY-female/GDX and XX-female/GDX respectively), the infusion of Ang II resulted in an increase in blood pressure and, with estradiol replacement, a depressor activational hormonal effect was also shown (panels C and D, respectively).

## DISCUSSION

Our results indicate that, in absence of activational hormonal effects, the interaction between the SCC and organizational hormonal action modulates the changes in arterial pressure due to acute Ang II infusion. Furthermore, estrogen and testosterone exert significant activational effects on changes in MAP during Ang II administration, but these responses are dependent on the SCC and/or the organizational hormonal backgrounds.

Studies analyzing activational hormonal effects on arterial blood pressure have demonstrated that treatment with estradiol attenuates hypertension in intact males and ovariectomized females, while administration of testosterone increases hypertension in gonadectomized males (Crofton and Share, 1997). Our data demonstrate that gonadectomized XY-male mice (without hormone replacement) showed no increase in mean arterial pressure due to Ang II infusion, but, when testosterone was replaced, XY-male/GDX+TP mice showed the expected increase in arterial pressure, clearly indicating the influence of activational testosterone. Furthermore, when estrogen was administered, no changes in blood pressure were observed compared to gonadectomized XY-male mice. Thus, our data demonstrate that, although E2 is largely considered to be protective in both male and female rodents, in XY-male gonadectomized mice, estradiol has no evident effect on blood pressure during acute Ang II infusion.

Moreover, gonadectomized XX-male, XX-female and XY-female mice (without hormone replacement) showed an increase in blood pressure in response to Ang II infusion and, when we performed β-estradiol hormone replacement, a depressor activational effect was observed in all cases, while no activational hormonal effect was observed for testosterone propionate. Previous studies have demonstrated that testosterone regulates (increases) the production of angiotensinogen, renin activity and the expression of the AT1-r receptor, favoring in this case the vasoconstrictor effect of RAS, and that these effects are significantly diminished after castration in males (Sampson et al., 2012). BP is significantly decreased in castrated male rats compared to intact control rats, and testosterone supplementation restored BP in testis-intact controls; furthermore, in female rats, dihydrotestosterone supplementation increased BP significantly compared to controls. Other studies demonstrated that testosterone downregulates angiotensin II type-2 receptor via the androgen receptor-mediated ERK1/2 MAP kinase pathway in rat aorta. AT2R mRNA and protein expression levels were lower in the aortas of males than females. In males, testosterone withdrawal by castration significantly elevated AT2R mRNA and protein levels, and testosterone replacement restored them. Moreover, in females, increasing androgen levels decreased AT2R mRNA and protein expression (Mishra et al., 2016).

Although testosterone is known to increase renin, ACE and AT1/AT2 balance, thus favoring the vasoconstrictor arm of the RAS, our results demonstrate that the pressor activational effect of testosterone is dependent not only on activational hormonal influence but also relies on the context and interaction of XY SCC and the organizational hormonal background. Our data shows that SCC has an important role in the pressor response to Ang II infusion under a masculine organizational background as, although both XY- and XX-male mice share the same organizational hormonal influence during critical periods of development, a clear difference was evident among XX- and XY-male mice in blood pressure response during ANG II infusion. While gonadectomized XY-male mice (without hormone replacement) showed no increase in mean arterial pressure due to Ang II infusion and, when testosterone was replaced, showed the expected increase in arterial pressure, XX-male mice showed the opposite effect. Gonadectomy without hormone replacement resulted in an increase in blood pressure in XX-male mice in response to Ang II infusion, while testosterone induced no activational effect. Furthermore, estradiol treatment prevented the increase in blood pressure in gonadectomized XX-male mice, while no activational β-estradiol effect was observed in XY-male/GDX-E2 mice, indicating an activational β-estradiol effect in male mice with the XX SCC background.

Our results also demonstrate an activational β-estradiol effect in XX- and XY-SCC females. Female mice (irrespective of having XX or XY sex chromosome complement but with the same organizational hormonal effect) as well as XX-male mice (with the typical XX-female genetic background) showed an attenuated blood pressure response when estradiol was replaced, while hormonal depletion by castration induced an increase in blood pressure under Ang II infusion. These results match those of Xue et al. (2007), who found that AngII-induced hypertension was greater in OVX and ERα KO female mice than in intact WT and OVX females treated with E2 for 21 days, and thus the latter revealed a downregulation of angiotensin-converting enzyme (ACE) transcript levels (Gallagher et al. 1999).

In female SHR rats in physiological estrus, microvessels were hyporeactive to Ang II in comparison to male SHR but, if females were ovariectomized, Ang II vasoconstriction was similar to that of males, and estrogen treatment abolished this difference. Silva-Antonialli et al. (2004) also demonstrated that the ARNm expression for AT1 was higher in aorta and mesenteric vessels of males than of females, that AT1mRNA expression in ovariectomized females was comparable to that of males, and that treatment with estrogen reversed the overexpression of AT1 receptor. Moreover, studies by Baiardi et al. (2005) demonstrated that estrogen upregulates renal AT2R. Thus, the lower AT1/AT2 ratio in intact females compared to males may at least partially explain the lower blood pressure observed in female SHR. In conjunction with this data, studies related to vascular function using the four core genotypes mouse model demonstrated that Ang II produced a greater contraction in the iliac artery in XY-male mice than in XX-female mice. The AT2R antagonist PD123319 revealed that this dilator response was mediated by the dilator AR2R only in intact XX-female mice (as no evident effect was observed in intact XX-sry male and XY-female mice), and that, when gonadectomized, the response could not be demonstrated, suggesting a significant interaction of the estradiol activational effect with the XX-SCC effect in the AT2-induced relaxation reported in intact XX-female mice (Pessoa et al., 2015).

Although previous studies by Ji et al. (2010) in gonadectomized mice of the FCG mouse model already demonstrated that chronic infusion of Ang II resulted, irrespective of sex, in a greater increase in blood pressure in XX-SCC mice than in XY-SCC mice, indicating the importance of SCC in the development of Ang II-hypertension in absence of activational hormonal effects, in the present study we evaluated the influence of the acute infusion of Ang II on blood pressure response, analyzing not only the effect of SCC and the organizational hormonal effect but also the activational hormonal influence of testosterone and estradiol in the sexually dimorphic blood pressure response.

In sum, this evidence isolates and highlights the contribution and interaction of SCC and the activational and/or organizational hormonal effects in sex differences in Ang II-blood pressure regulation. Our results indicate that, in the absence of activational hormonal effects, there is an interaction between the sex chromosome complement and organizational hormonal action, differentially modulating the changes in blood pressure due to an infusion of Ang II. Estrogen and testosterone exert important activational effects on changes in pressure during Ang II infusion; however, these effects are dependent on the sex chromosome complement and/or the organizational hormonal background.

## PERSPECTIVES

Clinical and experimental studies reveal that sex matters when it comes to blood pressure regulation, which highlights the importance of detailed study of the physiological differences among sexes in blood pressure regulation in physiological and pathophysiological states. Addressing in more detail the sources of physiological disparity between sexes and, in particular, the contribution of the SCC, organizational and activational hormonal effects involved in sex-related differences in RAS-blood pressure regulation, may help clarify not only the puzzling differences in how males and females regulate cardiovascular homeostasis, but also may have therapeutic importance and offer important insights into designing improved sex-tailored therapeutic treatments in the future.

## ABBREVIATIONS

AT1R: angiotensin II type-1 receptor
AT2R: angiotensin II type-2 receptor
ACE: angiotensin-converting enzyme
ACE2: angiotensin-converting enzyme type 2
E2: estradiol
GDX: gonadectomized
GDX group: gonadectomized mice without hormone replacement
GDX+E2: gonadectomized mice with β-estradiol replacement
GDX+TP: gonadectomized mice with testosterone propionate replacement
RAS: renin angiotensin system
SCC: sex chromosome complement
TP: testosterone propionate

## ACKNOWLEDGMENTS

We are grateful to Dr. Paul Burgoyne (Medical Research Council National Institute for Medical Research, London, UK) for providing the transgenic mice.

